## Supplementary Figures for "β-lactamase expression induces collateral sensitivity in *Escherichia coli*"

[1]*Servicio de Microbiología, Instituto Ramón y Cajal de Investigación Sanitaria (IRYCIS), Hospital Universitario Ramón y Cajal, Madrid, Spain.* [2]*Centro de Investigación Biomédica en Red de Enfermedades Infecciosas-CIBERINFEC, Instituto de Salud Carlos III, Madrid, Spain.* [3]*Centro Nacional de Biotecnología-CSIC, Madrid, Spain.* [4]*Centro de Investigación Biológica en Red de Epidemiología y Salud Pública-CIBERESP, Instituto de Salud Carlos III, Madrid, Spain*

All data supporting this article is provided as a *Supplementary Dataset* in a separate .csv file. Additionally, the following Supplementary Tables can be found as separate tabs in the *Supplementary Tables* file accompanying this article.

**Supplementary Table 1. Plasmids used in this study.**

**Supplementary Table 2. Antibiotic resistance and genomic features of ECOR strains.**

**Supplementary Table 3. Statistical analysis of ATLAs surveillance database.**

**Supplementary Table 4. Antibiotics used in this study.**

**Supplementary Table 5. Oligonucleotides used in this work.**

**Supplementary Figures**

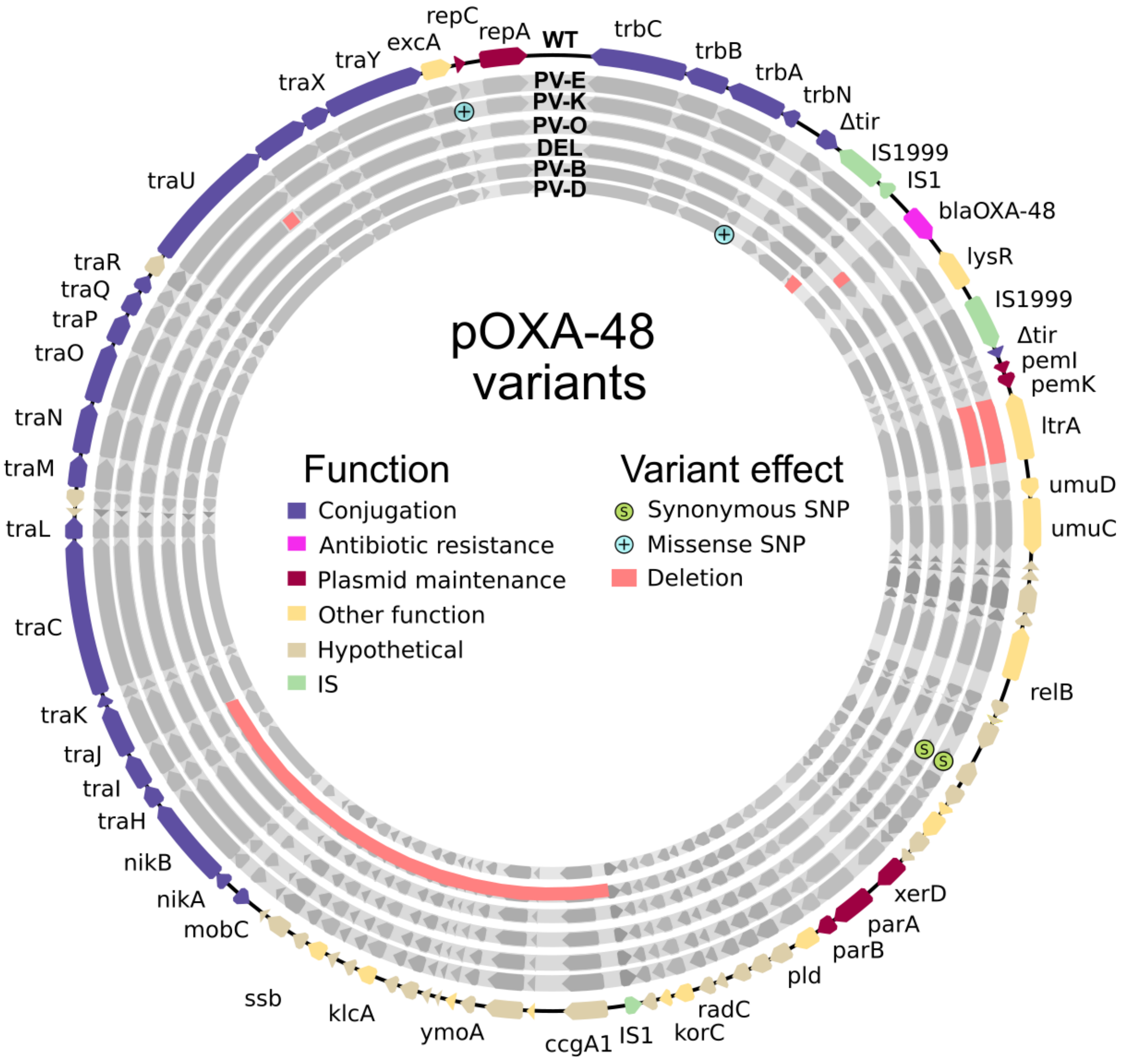


**Supplementary Figure 1. Representation of pOXA-48 variants used in this study**

The outer circle represents genes with known functions of the pOXA-48$_{\text{WT}}$ plasmid. Arrows indicate open reading frames coloured according to functional classification (see legend). Gene names are indicated in the outer circle. The inner circles represent the plasmid variants (PVs) used in this study. From outside to inside, pOXA-48$_{\text{PV-E}}$, pOXA-48$_{\text{PV-k,}}$ pOXA-48$_{\text{PV-O}}$, pOXA-48$_{\text{DEL,}}$ pOXA-48$_{\text{PV-B}}$, pOXA-48$_{\text{PV-D}}$. Point mutations are marked with blue or green circles for non-synonymous and synonymous mutations, respectively. Red tracks indicate deletions. See **Supplementary Table 1** for detailed information.

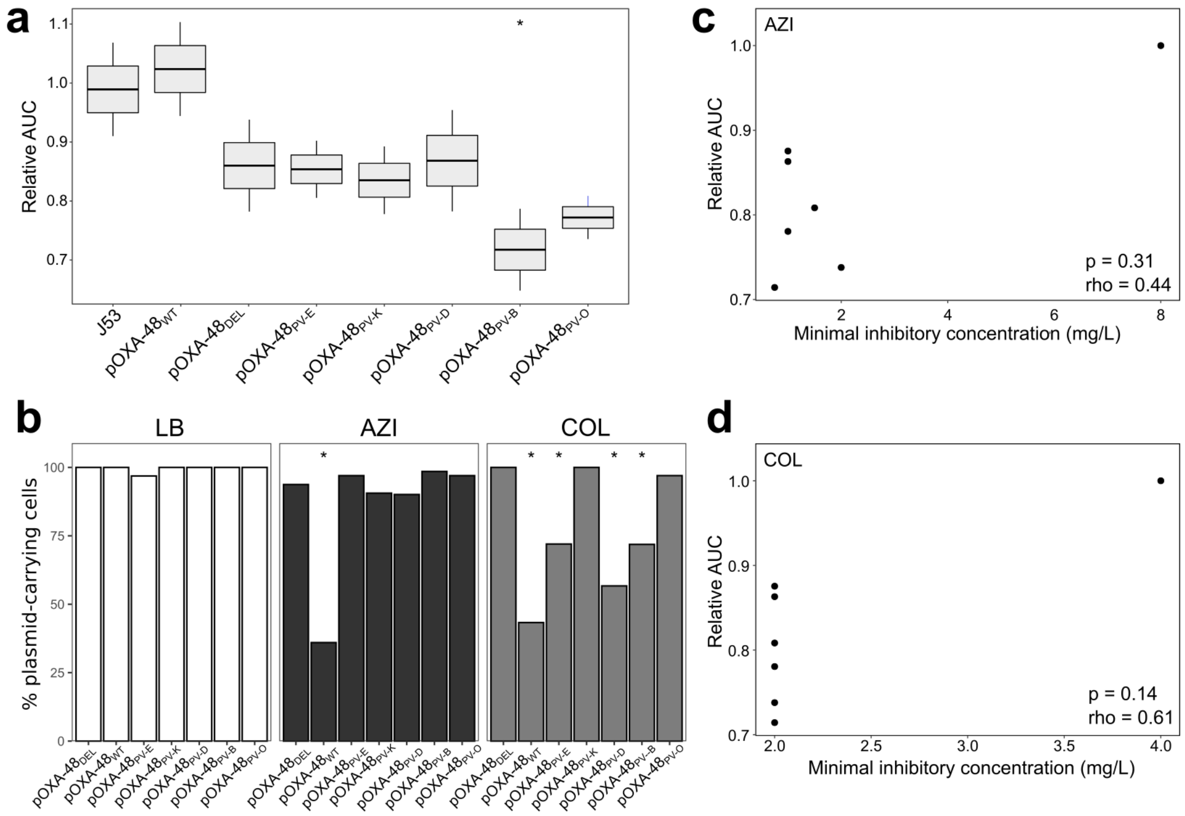


**Supplementary Figure 2. Analysis of growth and stability of plasmid-carrying bacterial population.**

**a)** Fitness (calculated as the relative area under the growth curve; AUC) of plasmid-carrying transconjugants relative to the plasmid-free J53 strain. Horizontal lines within boxes indicate median values, upper and lower hinges correspond to the 25th and 75th percentiles, and whiskers extend to observations within 1.5 times the interquartile range (six biological replicates). Asterisks indicate statistically significant differences (ANOVA Krustall-Wallis test $p < 0.05$). **b)** Plasmid stability of pOXA-48 variants after antibiotic treatment for 100 bacterial colonies as determined by PCR. In the absence of antibiotics, all populations maintained their respective pOXA-48 variant. Exposure to COL led to significant plasmid loss in 4 pOXA-48 variants (pOXA-$48_{WT}$, pOXA-$48_{PV\text{-}B}$, pOXA-$48_{PV\text{-}D}$, and pOXA-$48_{PV\text{-}E}$; Fisher's exact test $p<0.0001$ in all cases), and AZI treatment reduced the percentage of plasmid carriers in the pOXA-$48_{WT}$ variant (Fisher's exact test $p < 0.0001$). Notably, the pOXA-$48_{DEL}$ variant was stable across all conditions tested (Fisher's exact test $p > 0.2$), indicating that the loss of the pOXA-$48_{DEL}$ plasmid does not explain the abolition of the CS phenotype. **c)** and **d)** Lack of correlation between the fitness of plasmid-carrying clones (relative AUC) and the MIC (in mg/L) of azithromycin and colistin. Spearman's rho and p-values are indicated within each plot.

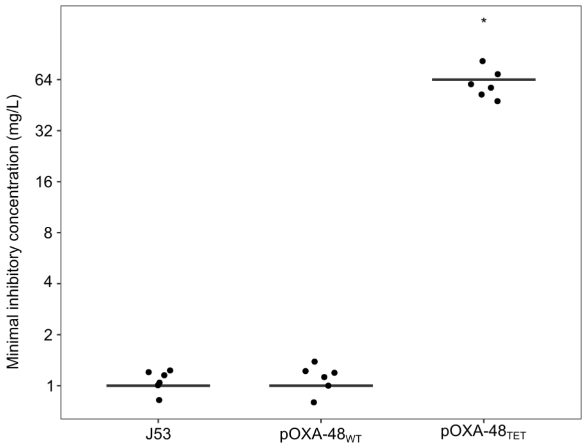


**Supplementary Figure 3. MIC of tetracycline for plasmid-free *E. coli* J53 and its derivatives carrying pOXA-48$_{WT}$ and pOXA-48$_{TET}$**

Horizontal lines indicate the median values of 6 biological replicates, indicated as individual data points. Asterisks denote statistically significant differences compared to plasmid-free J53 (Mann-Whitney u test $p < 0.0001$).

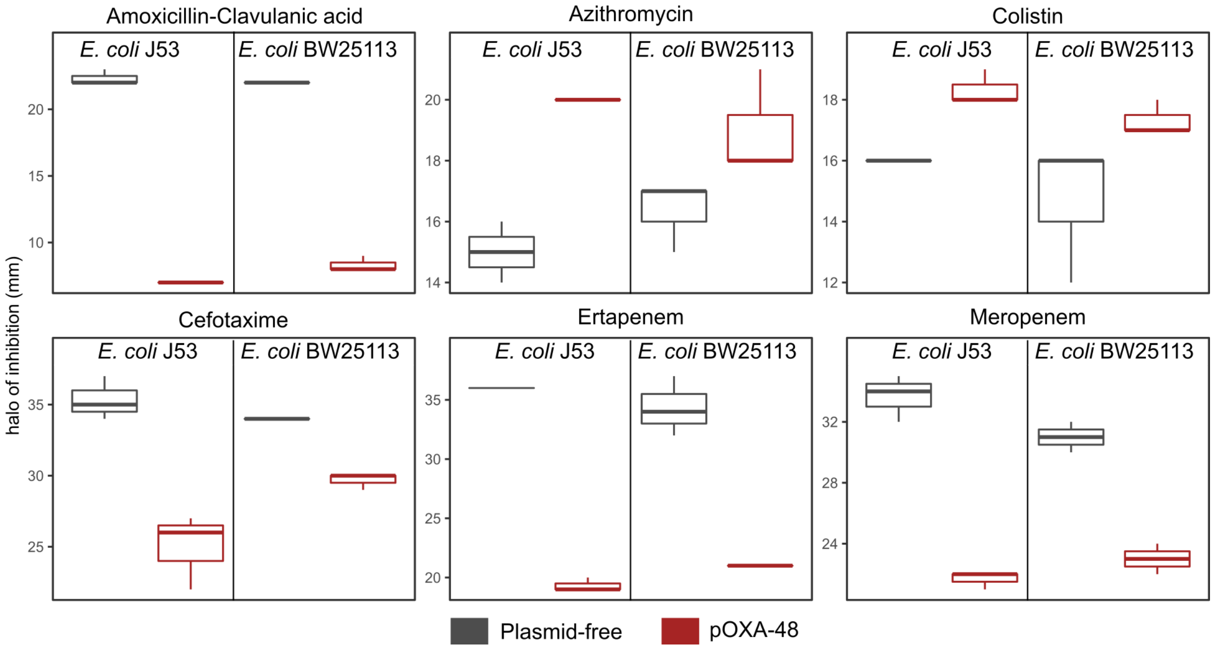


**Supplementary Figure 4. Comparison of antibiotic resistance profiles of *E. coli* J53 and BW25113.**

Boxplot representations of the inhibition halo diameters, in mm, obtained from disk-diffusion antibiograms of plasmid-free (grey) and plasmid-carrying (red) J53 or BW25113 strains. Horizontal lines within boxes indicate median values, upper and lower hinges correspond to the 25th and 75th percentiles, and whiskers extend to observations within 1.5 times the interquartile range.

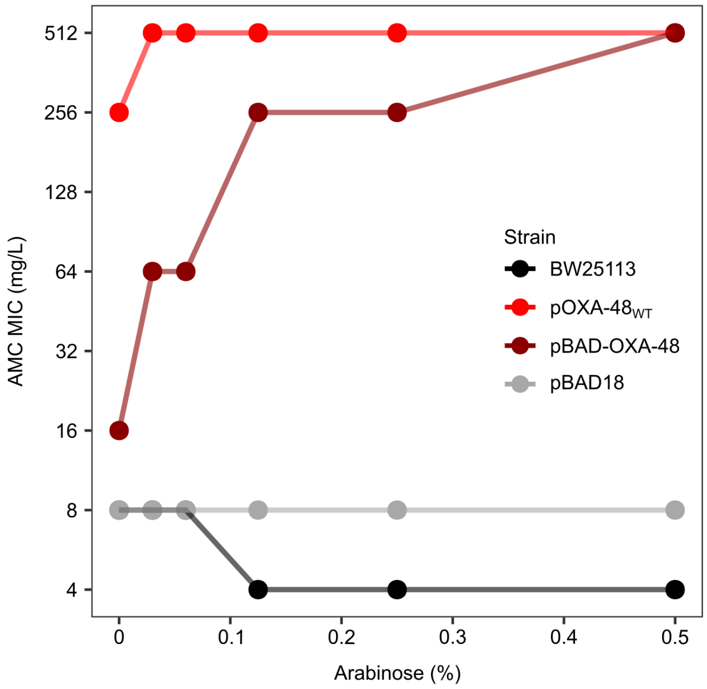


**Supplementary Figure 5. β-lactamase expression under the control of the $P_{BAD}$ promoter in the pBAD18 vector.**

The plot shows amoxicillin-clavulanic acid (AMC) resistance as a function of arabinose concentration for plasmid-free BW25113 strain (black), BW25113 carrying pOXA-$48_{WT}$ (light red) and BW25113 carrying the pBAD-OXA-48 (dark red) or the empty pBAD-18 plasmid (grey). Only at an arabinose concentration of 0.5 % (w/v) the strain carrying pBAD-OXA-48 shows resistance levels comparable to those provided by the pOXA-$48_{WT}$ plasmid. Data shows representative results from duplicate checkerboard assays.

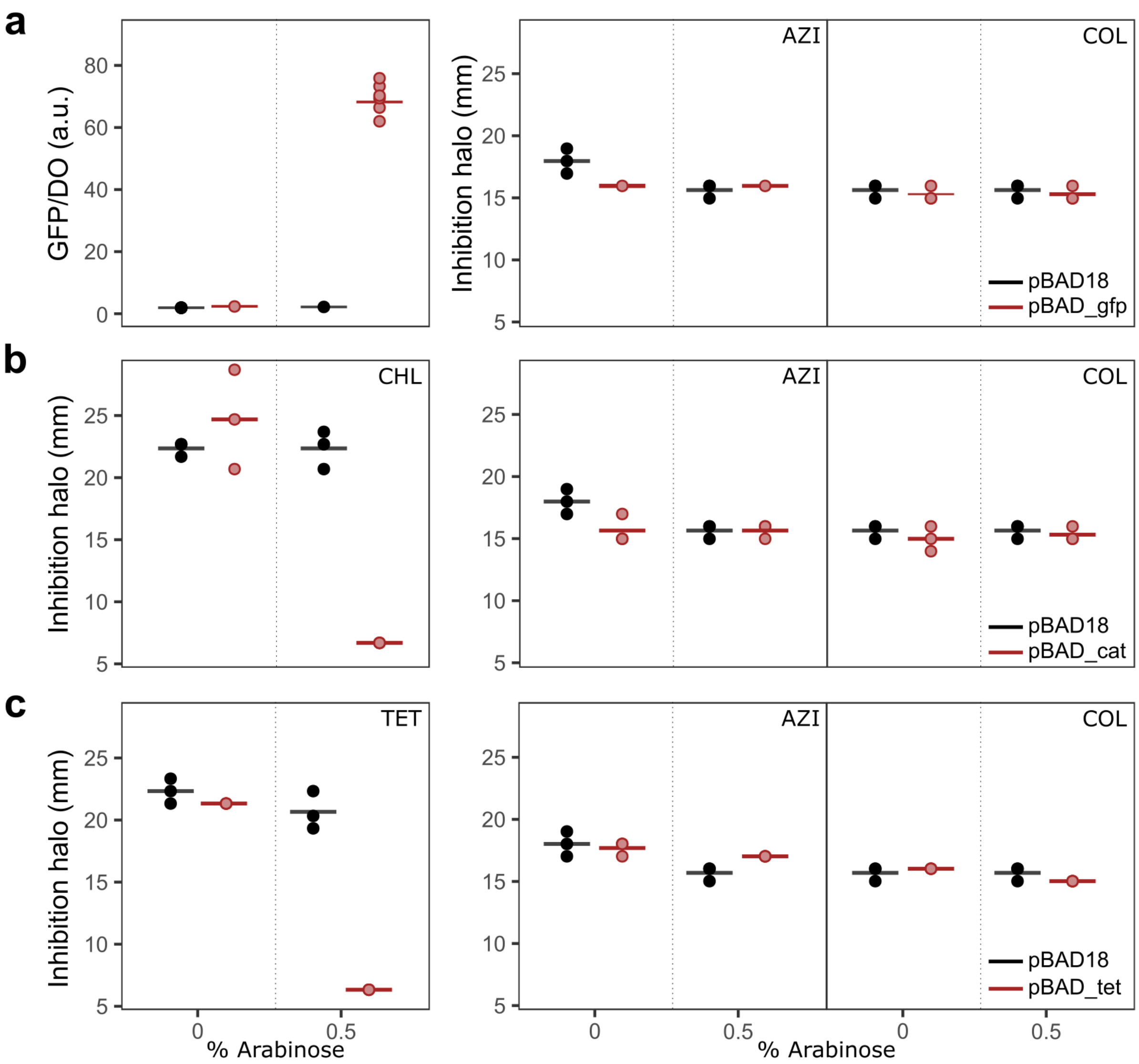


**Supplementary Figure 6. Expression of non-β-lactamase genes does not induce collateral sensitivity to AZI or COL**

**a)** Left: Fluorescence levels of the arabinose-induced pBAD-gfp construction and the empty pBAD18 plasmid. Right: AZI and COL inhibition halo diameters, in mm, obtained from disk-diffusion assays of strains carrying either the pBAD18 or the pBAD-gfp plasmid. Horizontal lines represent the median values, and the individual points represent independent replicates (n = 3). **b)** and **c)** Diameter of the inhibition halo, in mm, from disk-diffusion assays to different antibiotics of strains bearing the pBAD18, pBAD-cat (panel b) or pBAD-tet (panel c) plasmids. Horizontal lines represent median values and the individual points represent independent replicates (n = 3). The antibiotics used are indicated at the top right corner of each panel. AZI: azithromycin (15 mg); COL: colistin (10 mg); CHL: chloramphenicol (30 mg); TET: tetracycline (30 mg).

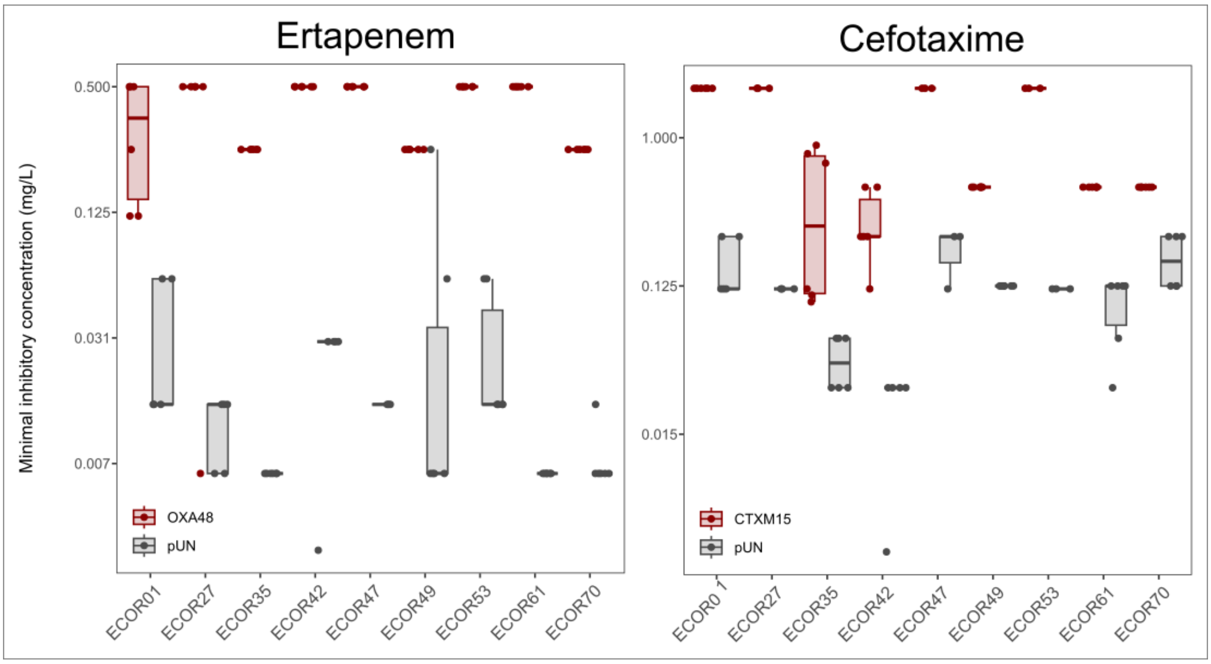


**Supplementary Figure 7. Antibiotic resistance profile of ECOR strains to ertapenem and cefotaxime.**

Boxplot representations of the MIC of ertapenem and cefotaxime, in mg/L, obtained from broth microdilution assays of the empty plasmid (pUN4) and β-lactamase-carrying plasmids ($bla_{OXA\text{-}48}$ and $bla_{CTX\text{-}M\text{-}15}$) in ECOR strains. Horizontal lines within boxes indicate median values, upper and lower hinges correspond to the 25th and 75th percentiles, and whiskers extend to observations within 1.5 times the interquartile range. Individual data points are also represented (6 biological replicates). The data points have been slightly jittered to avoid overlapping.

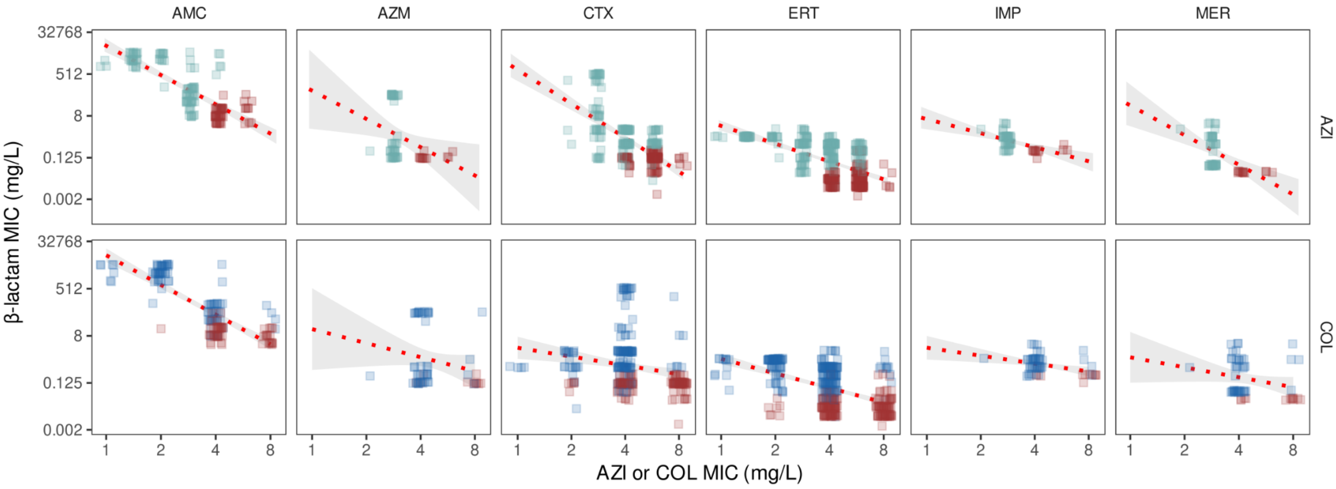


**Supplementary Figure 8. Experimental data shows negative correlations between β-lactam resistance and resistance to AZI (first row) and COL (second row).**

Blue points represent β-lactamase-free strains, and red points indicate the β-lactamase-carrying strains. Data points have been slightly jittered to avoid overlapping. The red dotted line represents the best linear fit for the data, and the grey-shaded zone covers the 95% confidence interval. All correlations are statistically significant (Spearman's correlation $p<0.04$). AMC: amoxicillin-clavulanic acid; ERT: ertapenem; CTX: cefotaxime; AZM: aztreonam; IMP: imipenem; MER: meropenem; AZI: azithromycin; COL: colistin.

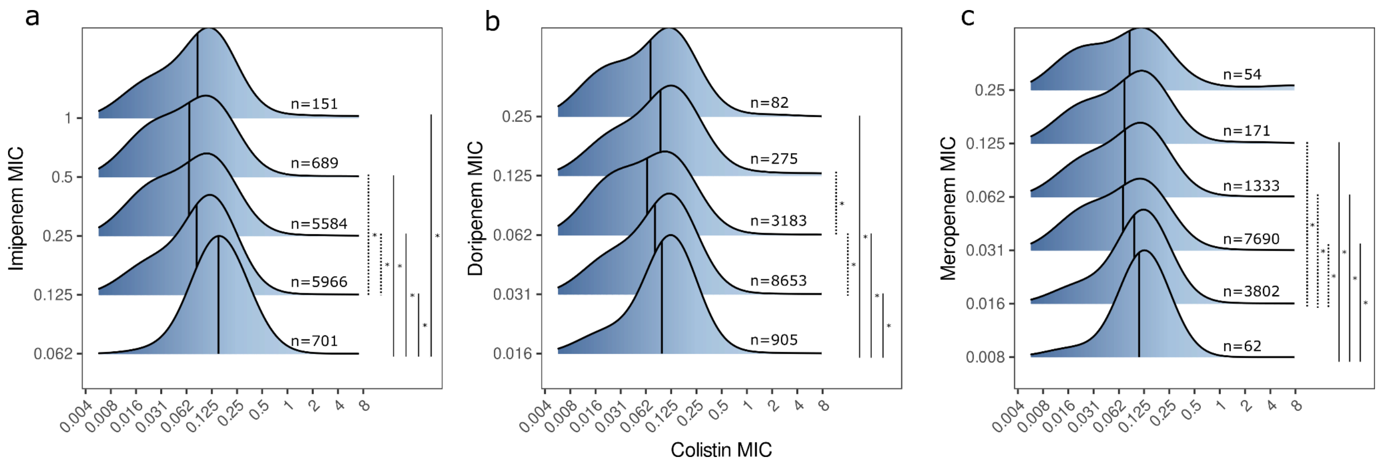


**Supplementary Figure 9. Carbapenem resistance negatively correlates with COL susceptibility.**

Colistin MIC distribution associated with each β-lactam MIC value, in mg/L, of **a)** imipenem, **b)** doripenem, and **c)** meropenem. The vertical line represented the mean colistin MIC of the distribution. Asterisks denote statistically significant differences between carbapenem concentrations (Dunn test after significant Kruskal-Wallis adjusted $p < 0.01$ in all cases).

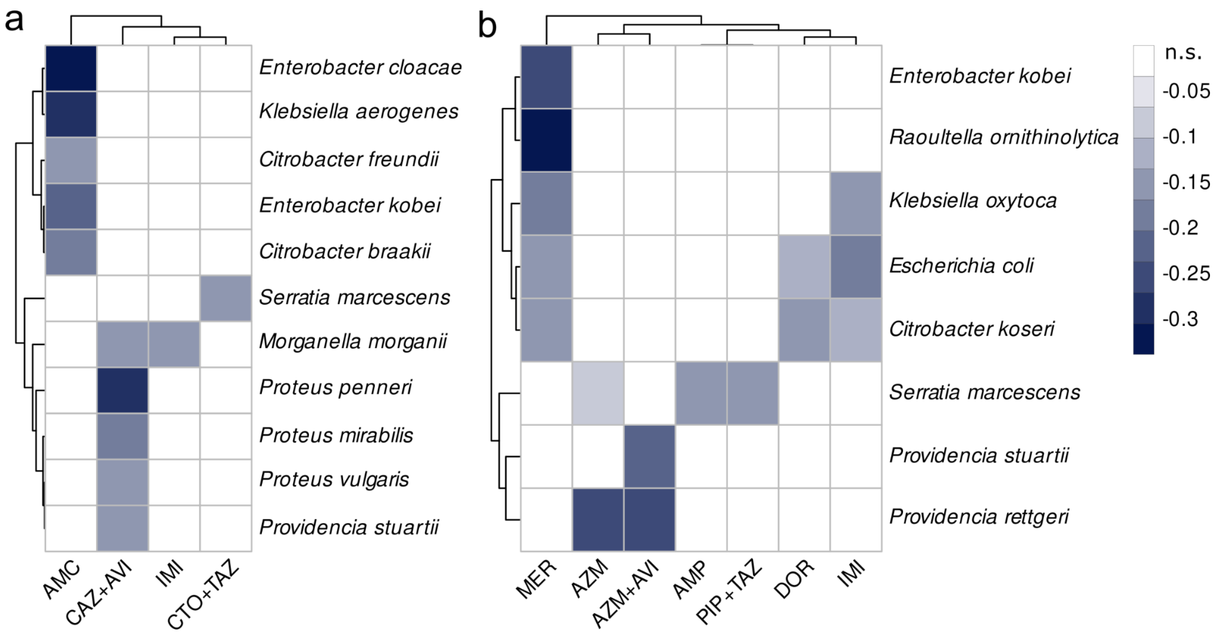


**Supplementary Figure 10. Heatmap revealing negative correlations between β-lactam resistance and COL (a) or COL + P-80 (b) in different species belonging to the Enterobacterales order (data from ATLAS).**

ATLAS contains susceptibility data for colistin and colistin supplemented with polysorbate 80 (P-80), a surfactant used to minimise non-specific colistin binding to plastic containers during broth microdilution experiments. Although the Clinical and Laboratory Standards Institute (CLSI) initially recommended using P-80[71], it was later discontinued in routine practice. Given that MIC values for both COL and COL + P-80 show good correlation in Gram-negative pathogens[72], in Fig. 5, we have decided to represent both COL and COL + P-80 data together. Antibiotic pairs with no correlation are depicted in white, and significant (Spearman's correlation $p < 0.05$) negative correlations between COL or COL + P-80 are shown in shades of blue. The strength of the colours is proportional to the strength of the correlation (Spearman's rho), as shown in the legend. AMC: amoxicillin-clavulanic acid, CAZ + AVI: ceftazidime + avibactam , IMI: imipenem, CTO + TAZ: ceftolozane + tazobactam, MER: meropenem, AZM: aztreonam, AZM + AVI: aztreonam + avibactam, AMP: ampicillin, PIP + TAZ: piperacillin + tazobactam, DOR: doripenem.
